# Nitrate availability modulates the temperature effect on N_2_O and N_2_ production from denitrification

**DOI:** 10.1101/2025.09.07.674766

**Authors:** Yueyue Si, Mark Trimmer

## Abstract

Nitrous oxide (N_2_O) can be both produced and subsequently reduced to dinitrogen gas (N_2_) via canonical denitrification, making the balance between these steps a key control on the net flux of this potent climate gas. Through a meta-analysis, we showed that net N_2_O and N_2_ production from denitrification respond differently to temperature, exhibiting distinct temperature sensitivities. In addition, nitrate availability plays a critical role in regulating this balance, yet only few studies have examined the combined effects of temperature and nitrate availability in natural sediments. Using ^15^N-isotope labelling and anoxic sediment incubations, we found that temperature effects on N_2_O and N_2_ production from denitrification were evident only under high nitrate levels (100 µM), while no significant temperature response occurred under low nitrate concentration (10 µM). At high nitrate availability, N_2_ production increased at higher temperatures, whereas net N_2_O production declined, leading to a lower production ratio of N_2_O to N_2_ at warmer temperatures. These findings suggest that in nitrogen-limited ecosystems, substrate availability plays a stronger role than temperature in regulating denitrification. More broadly, they provide insights into how nutrient loading and climate warming interact to shape nitrogen cycling and greenhouse gas emissions in aquatic ecosystems.

## Introduction

N_2_O has 273 times the warming potential of carbon dioxide (CO_2_) (Masson-Delmotte et al., 2021), is currently the primary driver of stratospheric ozone depletion (Ravishankara et al., 2009) and, most importantly, its atmospheric concentration has already risen by 25% since the 1850s (Meinshausen et al., 2011). Freshwater systems, especially small lakes and ponds, are increasingly recognised as significant sources of N_2_O, with their emissions increased by 126% over the same period (Li et al., 2024).

Microbial denitrification which proceeds via the pathway NO_3_^-^ →NO_2_^-^ →NO →N_2_O →N_2_ (Knowles, 1982), is a key process for both the production and reduction of N_2_O in freshwaters (Beaulieu et al., 2014; Beaulieu et al., 2011; Maavara et al., 2019), as well as in marine (Codispoti et al., 2001; Ji et al., 2018) and terrestrial ecosystems (Yu et al., 2023). As denitrification involves both the formation and consumption of N_2_O, its net emission to the atmosphere depends on the relative rates between these two steps. The activity of each step is regulated by environmental factors such as nitrate availability, dissolved oxygen concentration, and temperature (Codispoti et al., 2001; Ji et al., 2018; Kuypers et al., 2018).

Temperature is a key regulator of total N_2_O and N_2_ production via denitrification, with both generally increasing as temperature rises (Bailey and Beauchamp, 1973; Holtan-Hartwig et al., 2002; Keeney et al., 1979; Seitzinger et al., 1984; Silvennoinen et al., 2008). However, the effect of temperature on net N_2_O production – defined as total N_2_O production minus its consumption - varies widely among studies. Some studies report a positive relationship between net N_2_O production and temperature, suggesting that N_2_O production exceeds its reduction to N_2_ at higher temperatures (Dobbie and Smith, 2001; Duan et al., 2019; McKenney et al., 1984; Myrstener et al., 2016; Smith et al., 1998). Conversely, others observed a negative relationship, where N_2_ production increases more than N_2_O production as temperature rise, resulting in lower net N_2_O production (Silvennoinen et al., 2008). In addition, some studies find no significant relationship between temperature and net N_2_O production (Bailey, 1976; Del Prado et al., 2006; Lai and Denton, 2018). This variability underscores the complexity of temperature effects on denitrification and highlights the need for further investigation into how temperature interacts with other environmental drivers to regulate N_2_O dynamics and the balance in N_2_O:N_2_.

The availability of nitrate (NO_3_^-^) is another critical factor regulating the production of N_2_O and N_2_ via denitrification (Baulch et al., 2011; Beaulieu et al., 2011). However, many studies investigating the temperature dependence of N_2_O and N_2_ production have focused on nitrate-rich environments (Silvennoinen et al., 2008) or have experimentally added NO_3_^-^ concentrations far in excess of ambient concentrations (Jørgensen, 1989; Lai and Denton, 2018; Myrstener et al., 2016; Rysgaard et al., 2004). In some cases, sediments were subjected to prolonged exposure to high NO_3_^-^ enrichment for over three months, as in estuarine mesocosm studies, which significantly increased the temperature sensitivity of N_2_ production (Nowicki, 1994).

To date, studies examining the combined effects of nitrate availability and temperature on denitrification remain scarce (Lai and Denton, 2018; Myrstener et al., 2016). As a result, it is still unclear how the temperature sensitivity of N_2_O and N_2_ production varies across different nitrate concentrations. Most studies that have characterised the temperature sensitivities of both gases have focused on soils at typically high experimental NO_3_^-^ concentrations, whereas few have addressed aquatic sediments that are major sites of denitrification and nitrogen loss as N_2_O or N_2_ (Silvennoinen et al., 2008). Of the limited research in aquatic sediments, only two studies have characterised the temperature dependency of net N_2_O production from denitrification (Myrstener et al., 2016; Silvennoinen et al., 2008), and none has done so using direct measurements of ^15^N-labelled N_2_O. This is a critical gap, as measuring bulk N_2_O concentrations alone makes it difficult to disentangle the multiple microbial processes contributing to both N_2_O production and reduction (Zhu et al., 2025).

In this study, we first conducted a meta-analysis of existing data to predict the temperature sensitivities of N_2_O and N_2_ production, as well as their ratio, in N-rich soils and aquatic sediments. To assess how temperature and nitrate availability regulate N_2_O and N_2_ production in understudied N-limited aquatic sediments, we experimentally quantified the temperature dependencies of ^15^N-labelled N_2_O and N_2_ production from denitrification under varying ^15^N-nitrate concentrations. We addressed two key questions – (1) How does the net production of N_2_O and N_2_, and their ratio, respond to temperature variation in N-limited aquatic ecosystem? and (2) Does nitrate availability influence the temperature sensitivity of N_2_O and N_2_ production in such system?

## Methods

### Meta-analysis: Temperature sensitivities of net N_2_O and N_2_production from denitrification

To derive the temperature sensitivities of net N_2_O and N_2_ production from denitrification, and their ratios, here we compiled a list of studies that reported these rates and ratios under different temperatures in both aquatic sediments and terrestrial soils (Table 1). We fitted the net rate of N_2_O production, N_2_ production, and the N_2_O:N_2_ ratio into linear mixed-effect models to evaluate their temperature sensitivities (Bates et al., 2014). We then estimated the apparent activation energy (*Ē*) by fitting the natural log-transformed rate or ratio against the centred temperature term 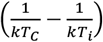 (Yvon-Durocher et al., 2014; Zhu et al., 2020):

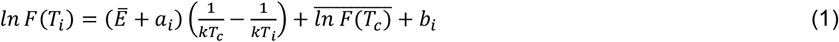

*k* is the Boltzmann constant (8.62 × 10^-5^ eV K^-1^, 1 eV = 96.485 kJ mol^-1^), while *T*_*c*_ was calculated by the sum of maximum and minimum of the inverted absolute temperature (in Kelvin) of the dataset, then divided by 2 (*T*_*c*_ = (max +min)/2). *T*_*i*_ is the absolute temperature from study *i* (*i* =1, 2, …,16). Further, to account for variances across the different studies, we included random slope (*a*_*i*_) and random intercept (*b*_*i*_) terms in the mixed-effects models.

**Table 1.** The studies used for the meta-analysis to characterise the temperature sensitivity of net N_2_O or N_2_production attibuted to denitrification in aquatic sediments and terrestrial soils. In most cases, denitrification was inferred rather than conclusively separated from other potential sources of N_2_O or N_2_. Inferred: N_2_O or N_2_ attributed to denitrification without confirmatory evidence to exclude other sources. Distinguished: N_2_O or N_2_ production from denitrification confimed by isotope-pairing techniques, which separated different sources. Number of measurements (*n*). In total, the dataset consisted of 920 measurements from 23 studies, i.e., 251 and 669 measurements in aquatic and terrestrial ecosystems, respectively. n.a.: not applicable.

| Ecosystem | Measurement | Denitrification | Substrate | Studies |
| --- | --- | --- | --- | --- |
| Aquatic | N<sub>2</sub>O | Inferred | NO<sub>3</sub><sup>-</sup>: 57 µM | (Myrstener et al., 2016) |
|  | N<sub>2</sub> | Inferred | Unspecified DIN | (Nowicki, 1994) |
|  | N<sub>2</sub> | Inferred | Ambient | (Seitzinger et al., 1984) |
|  | <sup>15</sup>N<sub>2</sub> | Distinguished | <sup>15</sup>NO<sub>3</sub><sup>-</sup>: 100 µM | (Brin et al., 2014; 2017) |
|  | <sup>15</sup>N<sub>2</sub> | Distinguished | <sup>15</sup>NO<sub>3</sub><sup>-</sup>: 50 µM | (Rysgaard et al., 2004) |
|  | <sup>15</sup>N<sub>2</sub> | Inferred | <sup>15</sup>NO<sub>3</sub><sup>-</sup>: 5.9 to 20.2 | (Veraart et al., 2011) |
|  | N<sub>2</sub>O, <sup>15</sup>N<sub>2</sub> | Inferred | NO<sub>3</sub><sup>-</sup>: 30 µM | (Silvennoinen et al., 2008) |
| Terrestrial | N<sub>2</sub>O | Inferred | NO<sub>3</sub><sup>-</sup>: 10204 µM | (Dobbie and Smith, 2001) |
|  | N<sub>2</sub>O | Inferred | NO<sub>3</sub><sup>-</sup>: 8571 µM | (Smith et al., 1998) |
|  | N<sub>2</sub>O | Inferred | NO<sub>3</sub><sup>-</sup>: 2536 µM | (Benoit et al., 2015) |
|  | N<sub>2</sub>O | Inferred | NO<sub>3</sub><sup>-</sup>: 10093 µM | (Del Prado et al., 2006) |
|  | N<sub>2</sub>O | Inferred | Ambient NO<sub>3</sub><sup>-</sup>: 286 µM | (McKenney et al., 1984) |
|  | N<sub>2</sub>O | Inferred | NO<sub>3</sub><sup>-</sup>: > 85 times ambient | (Kurganova and Lopes de Gerenyu, 2010) |
|  | <sup>15</sup>N<sub>2</sub>O | Inferred | <sup>15</sup>NO<sub>3</sub><sup>-</sup>: >18000 µM | (Duan et al., 2019) |
|  | N<sub>2</sub> | Inferred | NO<sub>3</sub><sup>-</sup>: 4 µM | (Qin et al., 2014) |
|  | N<sub>2</sub> | Inferred | NO<sub>3</sub><sup>-</sup>: 5497 µM | (Castaldi, 2000) |
|  | N<sub>2</sub> | Inferred | NO<sub>3</sub><sup>-</sup>: 4063 µM | (Holtan-Hartwig et al., 2002) |
|  | N<sub>2</sub>O, N<sub>2</sub> | Inferred | NO<sub>3</sub><sup>-</sup>: 4286 µM | (Bailey, 1976) |
|  | N<sub>2</sub>O, N<sub>2</sub> | Inferred | NO<sub>3</sub><sup>-</sup>: 8700 µM | (Keeney et al., 1979) |
|  | <sup>15</sup>N<sub>2</sub>O, <sup>15</sup>N<sub>2</sub> | Inferred | <sup>15</sup>NO<sub>3</sub><sup>-</sup>: >8000 µM | (Lai and Denton, 2018) |
|  | <sup>15</sup>N<sub>2</sub>O, <sup>15</sup>N<sub>2</sub> | Distinguished | <sup>15</sup>NO<sub>3</sub><sup>-</sup>: 8333 µM | (Yu et al., 2023) |
|  | N<sub>2</sub>O/N<sub>2</sub> | Inferred | Ambient NO<sub>3</sub><sup>-</sup>: 65 - 387 µM | (Maag and Vinther, 1996) |

### Optimisation of incubation conditions for characterising N_2_O and N_2_production from denitrification

Before characterising the temperature sensitivity of the gas products from denitrification, we carried out a trial experiment to determine the optimal substrate concentration and incubation time for the experiments with sediments collected from our well-established freshwater experimental ponds (Si et al., 2023).

Sediment cores were collected from the experimental ponds in December 2020, with each pond sampled at three different locations. Intact cores were transported to the laboratory in the dark on ice packs (4-hour trip) and then kept overnight at 4ºC. Before the experiment, the cores and pre-weighed vials (12 mL Exetainer, Labco) were put into an anaerobic glove box (5 ppm of residual O_2_, Belle Technology) which was constantly flushed with oxygen-free N_2_ (OFN) gas recycled through oxygen-scrubbing, catalytic cartridges. Anoxic medium was made by flushing N-free artificial pond water medium (Si et al., 2023) with OFN for 20 min. The top 2 cm of the cores for each pond were homogenized, then 2 mL of the sediment and 4 mL of the medium were added to each gas-tight vial to make a slurry. The slurry was added carefully without any sediment remaining inside the thread of the lid, as this may affect the sealing of the vial and lead to air contamination. The vials were then closed and pre-incubated in a temperature-controlled room (15ºC) for 16 hours to remove any residual porewater NO_*x*_^-^ and oxygen (Trimmer et al., 2013).

After the pre-incubation, vials were amended with 50 µL of different stocks of ^15^NO_3_^-^ (98% of ^15^N, Sigma Aldrich) to a final concentration range 10, 20, 50, or 100 µM, with un-amended vials as controls. Each concentration set was incubated for 0.5, 3, 6, 12, and 24 h (Table 2). Independent vials were used for each pond, substrate concentration, and time point. At each time point, the microbial activities in the vials were terminated by injecting 100 μL of formaldehyde (37 wt. %), with vials then equilibrated at room temperature (22ºC) until further analysis.

**Table 2.** Experiments designed to determine the optimal substrate and incubation length to characterise N_2_O and N_2_production from denitrification.

| Treatment | $^{15}\text{NO}_3^-$<br>($\mu\text{M}$) | Time<br>(h) | Targeted product(s) |
| --- | --- | --- | --- |
| Control | 0 | 0, 0.5, 3, 6, 12, 24 | $^{45}\text{N}_2\text{O}$, $^{46}\text{N}_2\text{O}$, $^{29}\text{N}_2$, $^{30}\text{N}_2$ |
| $^{15}\text{NO}_3^-$ | 10, 20, 50, 100 | 0, 0.5, 3, 6, 12, 24 | $^{45}\text{N}_2\text{O}$, $^{46}\text{N}_2\text{O}$, $^{29}\text{N}_2$, $^{30}\text{N}_2$ |

As the formaldehyde used to preserve the gas samples interferes with the colorimetric assay for NO_*x*_^-^ and NH_4_^+^ analyses, we prepared parallel samples for separate measurement of gases and nutrients. Samples for the nutrient measurement were immediately centrifuged and supernatants frozen at -20 °C at each time point until further analysis. To obtain the concentrations of porewater nutrients and the ^15^N-labelling of the NH_4_^+^ pool (FA), nutrient concentrations were measured by the automated wet-chemistry autoanalyzer (San^++^, SKALAR Analytical B.V.) with standard colorimetric techniques (Kirkwood, 1996). Calibration was performed against certified reference materials, traceable to NIST. The limits of detection were 0.05 μM, 0.1 μM and 0.2 μM for nitrite (NO_2_^*−*^), nitrite plus nitrate (NO_x_^*−*^: NO_2_^*−*^ + NO_3_^*−*^), and ammonium (NH_4_^+^), respectively.

For concentration of ^15^N-N_2_O, sub-samples from the headspace of each vial were transferred to an air-filled 12 mL gas-tight vial and measured on a continuous flow isotope ratio mass spectrometer (CF-IRMS, Delta V Plus, Thermo Finnigan) with an automated trace gas pre-concentrator (Precon, Thermo Finnigan). Calibration was performed with air, 0.12 ppm, 1.04 ppm, and a series of diluted 96 ppm N_2_O standards (BOC Limited), with a linear increase in the peak area over a range of 0.08 nmol to 5.85 nmol N_2_O in the vial. Concentrations of ^15^N_2_ were measured on the CF-IRMS in 100 µL of sample headspace. Drift in the signal of mass 30 was corrected by inserting air standards for every 10 samples (Si et al., 2023).

The net production of N_2_O and N_2_ in each vial was derived from the headspace concentration and the solubility of gases under the equilibrium temperature based on (Weiss and Price, 1980) for N_2_O and (Weiss, 1970) for N_2_ (Si et al., 2023). The production of ^15^N-N_2_O (^45^N_2_O + 2 × ^46^N_2_O) and ^15^N-N_2_ (^29^N_2_ + 2 × ^30^N_2_) was calculated from the excess gas production in the ^15^NO_3_^-^ treatments compared to that in the controls. After all the gas measurements, vials were centrifuged, supernatants removed and completely dried in the oven to obtain the dry weight for calculating weight-specific rates.

### Characterising the temperature sensitivities of N_2_O and N_2_production from denitrification

After we determined the optimal incubation conditions based on the trial experiment, we collected further sediment cores from eight experimental ponds in September 2021, to explore the temperature sensitivities of N_2_O and N_2_ production from denitrification.

As shown by the trial experiment, residual NO_3_^-^ remained in the incubation even after 16 hours of pre-incubation (Supplementary Fig. 1). Therefore, the vials were pre-incubated in a temperature-controlled room (15ºC) for a longer period (24 hours) to further remove any residual porewater NO_*x*_^-^. After the pre-incubation, vials were amended with different doses of ^15^NO_3_^-^ to a final concentration of 0 (controls), 10, or 100 µM (Table 3). Each concentration set was incubated for 3 hours, which is the optimal time determined from the trial experiment (Supplementary Fig. 2). Other details for sediment collection, sample preparations and measurements remained the same as in the trial experiments.

**Table 3.** Experiments designed to characterise the temperature sensitivities of N_2_O and N_2_ production via denitrification, and also test for any anammox potential, and N_2_O production from nitrification. As both denitrification and anammox produce N_2_, isotopic labelling with different ^15^N-substrates was used to distinguish between the two processes. Anammox activity can be confirmed if excess ^29^N_2_ is produced in the ^15^NH_4_^+^+^14^NO_3_^-^ treatment relative to ^15^NH_4_^+^-only treatment. All treatments were applied to independent sediment samples collected from eight ponds.

| Treatment | <sup>15</sup>NO<sub>3</sub><sup>-</sup><br>(μM) | <sup>15</sup>NH<sub>4</sub><sup>+</sup><br>(μM) | Temperature<br>(°C) | Targeted product(s) |
| --- | --- | --- | --- | --- |
| **Denitrification** |  |  |  |  |
| Control | 0 | n.a. | 5, 10, 15, 20, 25 | <sup>45</sup>N<sub>2</sub>O, <sup>46</sup>N<sub>2</sub>O, <sup>29</sup>N<sub>2</sub>, <sup>30</sup>N<sub>2</sub> |
| <sup>15</sup>NO<sub>3</sub><sup>-</sup> | 10 or 100 | n.a. | 5, 10, 15, 20, 25 | <sup>45</sup>N<sub>2</sub>O, <sup>46</sup>N<sub>2</sub>O, <sup>29</sup>N<sub>2</sub>, <sup>30</sup>N<sub>2</sub> |
| **Anammox** |  |  |  |  |
| <sup>15</sup>NH<sub>4</sub><sup>+</sup> | n.a. | 10 or 60 | 15 | <sup>29</sup>N<sub>2</sub> |
| <sup>15</sup>NH<sub>4</sub><sup>+</sup>+<sup>14</sup>NO<sub>3</sub><sup>-</sup> | n.a. | 10 or 60 | 15 | <sup>29</sup>N<sub>2</sub> |
| **Nitrification** |  |  |  |  |
| Control | n.a. | 0 | 15 | <sup>45</sup>N<sub>2</sub>O, <sup>46</sup>N<sub>2</sub>O |
| <sup>15</sup>NH<sub>4</sub><sup>+</sup> | n.a. | 22 or 44 | 15 | <sup>45</sup>N<sub>2</sub>O, <sup>46</sup>N<sub>2</sub>O |
| <sup>15</sup>NH<sub>4</sub><sup>+</sup>+ATU | n.a. | 44 | 15 | <sup>45</sup>N<sub>2</sub>O, <sup>46</sup>N<sub>2</sub>O |

### Characterising the potential of anammox and nitrification

As both denitrification and anammox (NH_4_^+^ + NO_2_^-^ →N_2_) produce N_2_ (Dalsgaard et al., 2003; Kuypers et al., 2003; Trimmer et al., 2013), distinguishing between these two processes is critical. This can be achieved by analysing the production of ^29^N_2_ (^14^N^15^N) and ^30^N_2_ (^15^N^15^N) from different combinations of ^15^N-labeled substrates (Brin et al., 2014; Dalsgaard et al., 2003; Kuypers et al., 2003; Trimmer et al., 2013). According to isotope-pairing principles, the presence of anammox is indicated by the production of excess ^29^N_2_ production in a treatment with ^14^NO_3_^-^ and ^15^NH_4_^+^, relative to a treatment with ^15^NH_4_^+^ alone - since anammox couples one nitrogen atom from ^14^NO_3_^-^ and one from ^15^NH_4_^+^ to form N_2_ (Dalsgaard et al., 2003; Trimmer et al., 2013). To test for the potential of the anammox reaction, two additional sets of incubations were performed using the same sediment samples as in the denitrification experiments: a ^15^NH_4_^+^ treatment, and a ^15^NH_4_^+^ plus ^14^NO_3_^-^ treatment (Table 3). Each treatment included two concentrations of ^15^NH_4_^+^ (final concentration of 10 or 60 µM, prepared from 98% ^15^N-NH_4_^+^, Sigma Aldrich) to test for anammox activity under both low and high substrate availability. In the combined treatment, ^14^NO_3_^-^ was added at a final concentration of 100 µM, consistent with the high ^15^NO_3_^-^ concentration used in the denitrification experiment above. After incubations, ^29^N_2_ concentrations were measured in each treatment to assess anammox activity.

In addition, we also tested the potential of N_2_O production from nitrification (NH_4_^+^ →NH_2_OH →N_2_O). Sediment cores were collected from eight ponds in February 2022, with the same sampling techniques as used in the denitrification experiments. In the laboratory, the top 2 cm of sediments for each pond were homogenised, then 2 or 3 mL of the sediment and 2.7 mL of artificial pond medium were added to each 12 mL vial to make an oxic slurry. Vials were amended with ^15^NH_4_^+^ (final concentration 22 µM or 44 µM) with or without allylthiourea (ATU) (100 µL from 2.8 mM stock, final concentration ∼80 µM), where ATU was used to block the oxidation of NH_4_^+^ (Ginestet et al., 1998). With 44.2 µM of ^15^NH_4_^+^ added, the ^15^N-labelling of NH_4_^+^ (F_A_) in the treatment was 0.85 ± 0.07, on average (Supplementary Table 1). Un- amended vials were prepared as controls to account for any activity of nitrification from the background NH_4_^+^ (Table 3). Independent vials were used for each pond and treatment and incubated for 0, 3, 8, 18, or 24 hours. All vials were incubated at 15ºC, which was the annual average temperature in the ambient ponds (Si et al., 2023). The microbial activities in samples were terminated by formaldehyde at each time point, with N_2_O measured with the same methods as used in the denitrification experiments.

### Statistical analysis

To derive the temperature sensitivities of net N_2_O and N_2_ production, and the N_2_O:N_2_ ratio from denitrification in the N-limited ponds, we estimated the apparent activation energy by fitting the natural log-transformed rate of net production of N_2_O or N_2_ against the centred temperature term 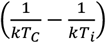 using equation (1), except that *i* (*i* =1, 2, …,8) now denotes the different ponds and *T*_*C*_ denotes the average incubation temperature (15°C).

Statistical analysis and plotting were performed in R version 4.5.0 (Team, 2021) using RStudio. For meta-analysis, as the studies used different normalisation methods, we standardised the data by subtracting the study-specific intercept from the rate of N_2_O or N_2_ production for each study (Yvon-Durocher et al., 2014; Zhu et al., 2020). For our incubations with pond sediments, we standardised the data by subtracting the pond-specific intercept from the net production rate of N_2_O or N_2_ for each pond. Models were ranked by the small sample-size corrected Akaike Information Criterion (AICc) using the ‘MuMIn’ package. The best-fitting model was determined by the lowest AICc score and the activation energies (Ē), in the unit of eV, derived from the slope of the best-fitting models.

## Results

### Meta-analysis

#### Temperature sensitivities of N_2_O and N_2_production from denitrification

From the meta-data analysis of previous studies, net N_2_O production from denitrification increased at higher temperatures, with an apparent activation energy (Ea) of 0.41 eV (95% CI of 0.29 to 0.54 eV, Fig. 1**a**). The temperature response of net N_2_O production was consistent across aquatic sediments and terrestrial soils (Supplementary Table 2, likelihood ratio test comparing M0 and M2 for net N_2_O, *p* > 0.05), although data from aquatic sediments remain limited. Similarly, N_2_ production from denitrification was higher at warmer temperatures, with an activation energy at 0.6 eV (95% CI of 0.45 to 0.75 eV, Fig. 1; and Supplementary Table 2 comparing best-fitting model M0 and null model, *p* < 0.001). The temperature sensitivity of N_2_ production was also consistent between aquatic and terrestrial ecosystems (Supplementary Table 2, likelihood ratio test comparing M0 and M2 for N_2_, *p* > 0.05).

**Fig. 1.**
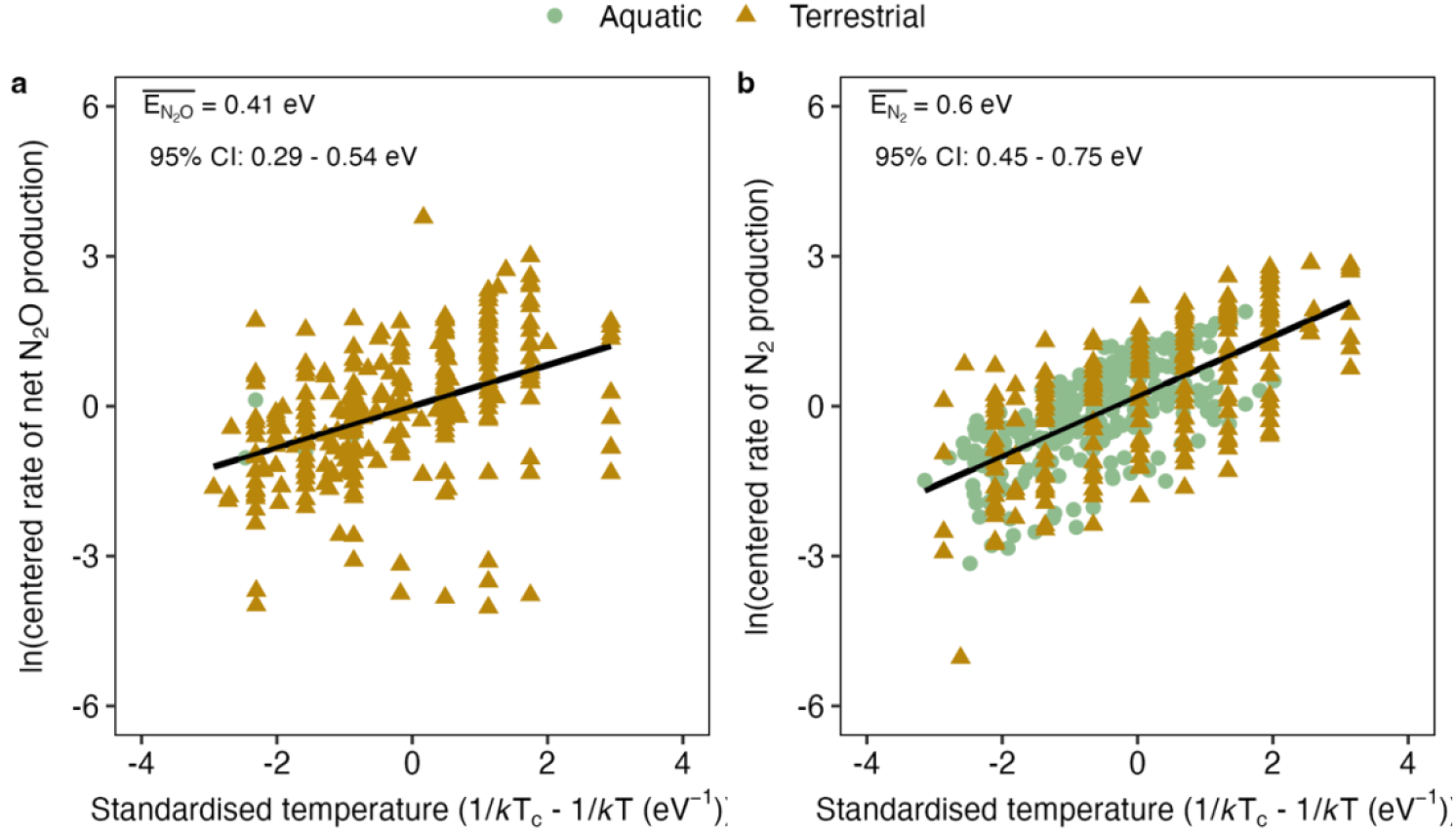
Meta-analysis fitting published rates of net N_2_O and N_2_production inferred from denitrification as a function of temperature with both aquatic sediments and terrestrial soils. The rate of net N_2_O and N_2_ production increased at higher temperatures in both aquatic sediments and soils. We visualised the data using the “Visreg” package in R (Breheny and Burchett, 2017) plotting the data as the partial residuals (brown and green circles) from the best fitting models after the random effects were accounted for and with the overall estimate for temperature sensitivity given as black lines (Supplementary Table 2) . The inverted absolute temperature was centered as Tc = (max+min)/2, while the rate of N_2_O or N_2_ production was natural log (ln) transformed and then centered by subtracting each study-specific intercept. For net N_2_O production, *n* = 8 and 265 measurements in aquatic and terrestrial, respectively, whereas for N_2_ productions, *n* = 239 and 208 measurements in aquatic and terrestrial, respectively. Because normalisation methods differed among studies, we standardised the data by centring each study’s N_2_O or N_2_ production rates on its specific intercept.

From studies reporting parallel measurements of net N_2_O and N_2_ production, we derived the temperature sensitivity of the N_2_O:N_2_ ratio, which showed a negative relationship with temperature (Fig. 2). We also predicted the temperature sensitivity of the N_2_O:N_2_ ratio using the independent datasets of net N_2_O and N_2_ shown in Fig. 1. The activation energy of the N_2_O:N_2_ ratio derived from the two approaches did not differ statistically (*χ*^*2*^ = 1.46, *p* = 0.23; linear hypothesis test for fixed effects using the ‘car’ package in R). These results showed that while both net N_2_O and N_2_ production from denitrification increased with temperature, their differing temperature sensitivities led to a negative relationship between the N_2_O:N_2_ ratio and temperature.

**Fig. 2.**
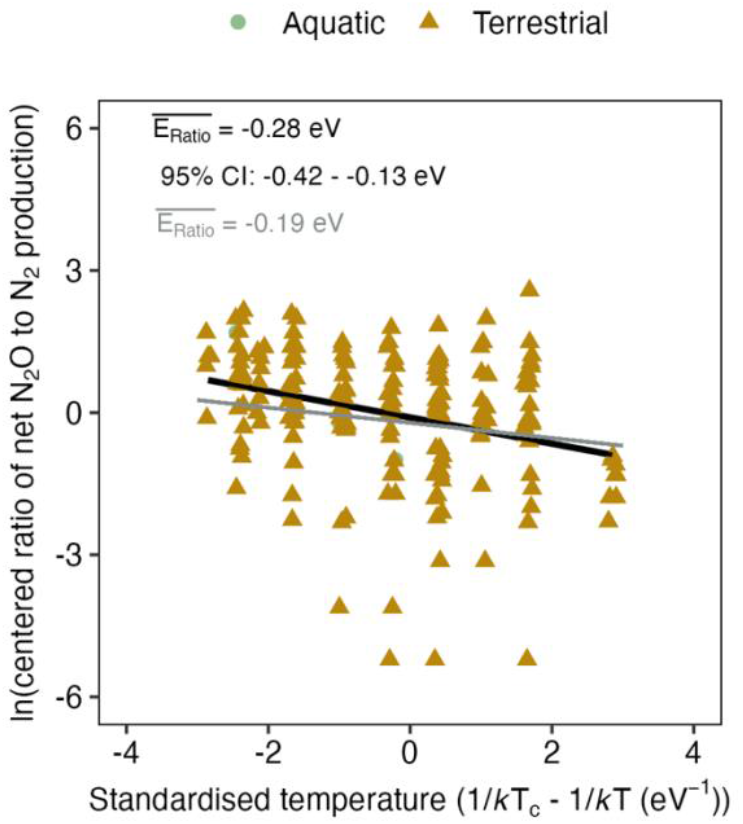
Ratio of net N_2_O to N_2_production from denitrification as a function of temperature, based on incubations from both aquatic sediments and terrestrial soils. *n* = 4 and 196 measurements for aquatic sediments and terrestrial soils, respectively. The black line gives the activation energy derived from the slope of the best fitting linear mixed-effects model for actual parallel measurements of net N_2_O and N_2_ production (Supplementary Table 2, Model M0 for N_2_O:N_2_). The grey line shows the activation energy of the N_2_O:N_2_ ratio, predicted from the independent net N_2_O and N_2_ datasets shown in Fig. 1. The two estimates did not differ significantly (*χ*^*2*^ = 1.46, *p* = 0.23; linear hypothesis test for fixed effects using the ‘car’ package in R).

However, while these previous studies commonly inferred denitrification as the source of the measured N_2_O or N_2_ production, most did not explicitly distinguish it from other microbial pathways contributing to N_2_O or N_2_ production (Table 1). As a result, the reported temperature sensitivities may reflect a composite signal from multiple processes, rather than denitrification alone.

#### Optimal incubation conditions for characterising net production of N_2_O and N_2_from denitrification

No excess production of ^29^N_2_ was detected in either the ^15^NH_4_^+^ only treatments or those amended with ^14^NO_3_^-^ compared to the controls (Supplementary Fig. 3). Likewise, adding extra ^14^NO_3_^-^ to the ^15^NH_4_^+^ treatments did not increase ^29^N_2_ concentrations (*p* = 0.61, Two-Sample *t*-test), confirming the absence of anammox. This agrees with previous studies showing no anammox activity in these pond sediments across three seasons (Warren, 2017). Nitrification was also negligible as a source of N_2_O, detected only in one of eight ponds and at much lower rates than denitrification (Supplementary Fig. 4). These results indicate that denitrification was the dominant pathway responsible for both N_2_O and N_2_ production in the studied ponds.

In the trial experiments, ^15^N-N_2_O production peaked and was subsequently reduced during all incubations (Supplementary Fig. 2**a**). At ^15^NO_3_^-^ concentrations below 100 µM, ^15^N-N_2_O reached its peak within ∼0.5 h of incubation and declined to zero before 24 h in most incubations. In contrast, with 100 µM ^15^NO_3_^-^, the ^15^N-N_2_O peak occurred later (at ∼3 h) and reached higher production than in the lower-concentration treatments. Meanwhile, ^15^N-N_2_ accumulated continuously over 24 h in incubations with ^15^NO_3_^-^ additions higher than 10 µM, but at 10 µM of ^15^NO_3_^-^ it plateaued after ∼12 h (Supplementary Fig. 2**b**).

The availability of nitrate influenced both the magnitude and the ration of N2O to N2?. Production rates of both ^15^N-N_2_O and N_2_ increased with increasing ^15^NO_3_^-^ concentrations (Fig. 3**a**, 3**b**). Specifically, ^15^N-N_2_O production showed a near-linear increase across the full concentration range (0 -100 µM), reaching a maximum of 223.27 nmol g^-1^ h^-1^ at 100 µM ^15^NO_3_^-^ (Fig. 3**a**). ^15^N-N_2_ production also increased with nitrate, but the response was gentler (Fig. 3**b**). The ratio of ^15^N-N_2_O to ^15^N-N_2_ production increased up to 50 µM) and then plateaued (Fig. 3**c**).

**Fig. 3.**
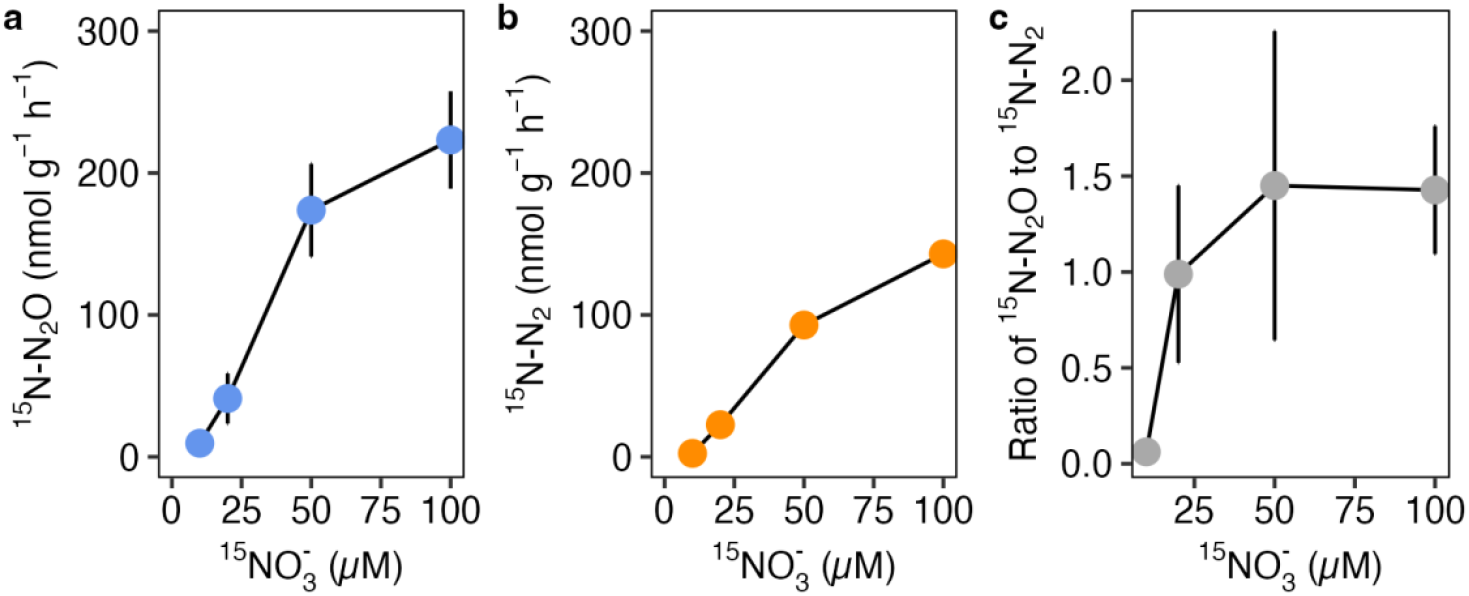
Rate of^15^N_2_O and^15^N_2_production and their ratios, from independent sediment incubations amended with varying concentrations of^15^NO_3_^-^. **a**, Rate of ^15^N_2_O production calculated from the 3-h incubation. **b**, Rate of ^15^N_2_ production calculated from the 24-hour incubation. Both gases showed increased production with higher ^15^NO_3_^-^ availability, with the rate of increase plateaued at 100 µM of ^15^NO_3_^-^. **c**, Ratio of ^15^N_2_O to ^15^N_2_ production across ^15^NO_3_^-^ concentrations. Dots in each plot represent means, with error bars indicating standard error (*n* = 6 ponds per concentration of ^15^NO_3_^-^ at each time point).

In addition, 43% of the ^15^NO_3_^-^ added was reduced to ^15^N_2_ after 24 h of incubation, on average (Supplementary Fig. 5), which was consistent across different ^15^NO_3_^-^ concentrations (*p* = 0.52, df = 14, Kruskal-Wallis test). This indicates that the fraction of the NO_3_^-^ reduced to the end-product N_2_ was not affected by NO_3_^-^ availability over the tested range of 10 to 100 µM.

#### Temperature sensitivities of N_2_O and N_2_ net production from denitrification

Based on results from the trial experiments, sediments were incubated with 100 µM of ^15^NO_3_^-^ for approximately 3 h to characterise the temperature sensitivity of both the production of N_2_O and N_2_. In addition, as the concentration of NO_3_^-^ in the ponds was low (< 2 µM, Supplementary Table 3), additional samples were incubated with a lower concentration of ^15^NO_3_^-^ (10 µM) to characterise the effect of lower substrate availability on the denitrification gas products.

After the 3-hour incubation, net production of both ^15^N_2_O and ^15^N_2_ was detectable in the large majority of incubations (95%, 152 out of 160 incubations) with either 10 µM or 100 µM of ^15^NO_3_^-^ added. Production of N_2_O and N_2_ responded differently to temperature depending on the availability (concentration) of NO_3_^-^. At 10 µM ^15^NO_3_^-^ net production of both ^15^N_2_O and ^15^N_2_ did not increase significantly at higher incubation temperatures (Fig. 4**a**, 4**c, M10.a** compared to M10.b for both N_2_O and N_2_, Supplementary Table 4). In contrast, with 100 µM ^15^NO_3_^-^, net production of both ^15^N_2_O and ^15^N_2_ was sensitive to increasing temperature (*p* < 0.001, **M100.a** compared to M100.b for both N_2_O and N_2_, Supplementary Table 4), but with opposite temperature sensitivities. That is, net N2O production decreased at -0.26 eV while N2 production increased at 0.43 eV (Fig. 4**b**, 4**d**). Between temperatures of 5ºC to 25ºC, ^15^N_2_O decreased, while ^15^N_2_ production increased at higher temperatures.

**Fig. 4.**
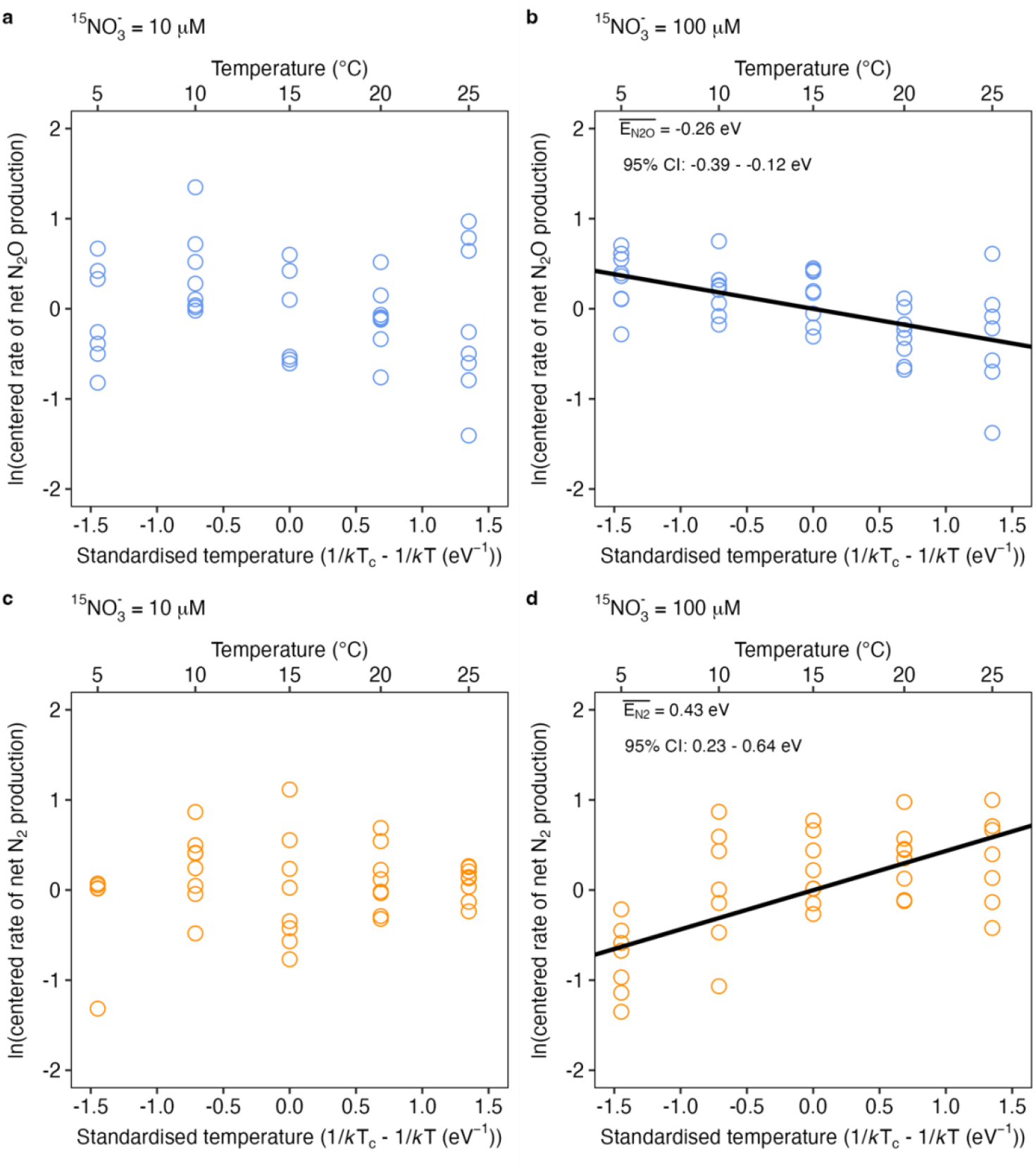
Temperature sensitivities of the net production rate (nmol g^-1^) of N_2_O and N_2_from denitrification under nitrate-limited or nitrate-replete conditions. **a**, Net production rate of ^15^N_2_O was consistent between different temperatures with 10 µM ^15^NO_3_^-^ added. **b**, ^15^N_2_O decreased at higher temperatures with 100 µM of ^15^NO_3_^-^. **c**, Net production of ^15^N_2_ was consistent between different temperatures with 10 µM ^15^NO_3_^-^ added. **d**, ^15^N_2_ accumulated at higher temperatures with 100 µM of ^15^NO_3_^-^. The temperature was centered at the median temperature of all the data points, i.e. 15°C, while the net production rates of N_2_O and N_2_ were natural log (ln) transformed and then centered by subtracting the pond-specific intercepts. we visualized the data using the “Visreg” package in R (Breheny and Burchett, 2017), with the lines in **b** and **d** showing the best-fitting linear mixed-effect model (Supplementary Table 4). The data shown are with the full range of incubation temperatures from 5ºC to 25ºC. *n* = 37, 39, 38, and 38 incubations, respectively, for panels **a** - **d**, conducted using sediments from 8 different ponds for each ^15^NO_3_^-^ treatment.

Furthermore, at 10 µM ^15^NO_3_^-^, the ^15^N_2_O:^15^N_2_ ratio was consistent across temperatures (Fig. 5**a**, *p* > 0.05, **M10.a** compared to M10.b for the ratio, Supplementary Table 4), whereas at100 µM ^15^NO_3_^-, 15^N_2_O:^15^N_2_ decreased exponentially from 5ºC to 20ºC (Fig. 5**b**, *p* < 0.001, **M100.a** compared to M100.b for the ratio, Supplementary Table 4) at -0.7 eV (95% CI: -0.93 to -0.47 eV, Fig. 5**b**) leading to a higher accumulation of N_2_O relative to N_2_ from denitrification at colder temperatures.

**Fig. 5.**
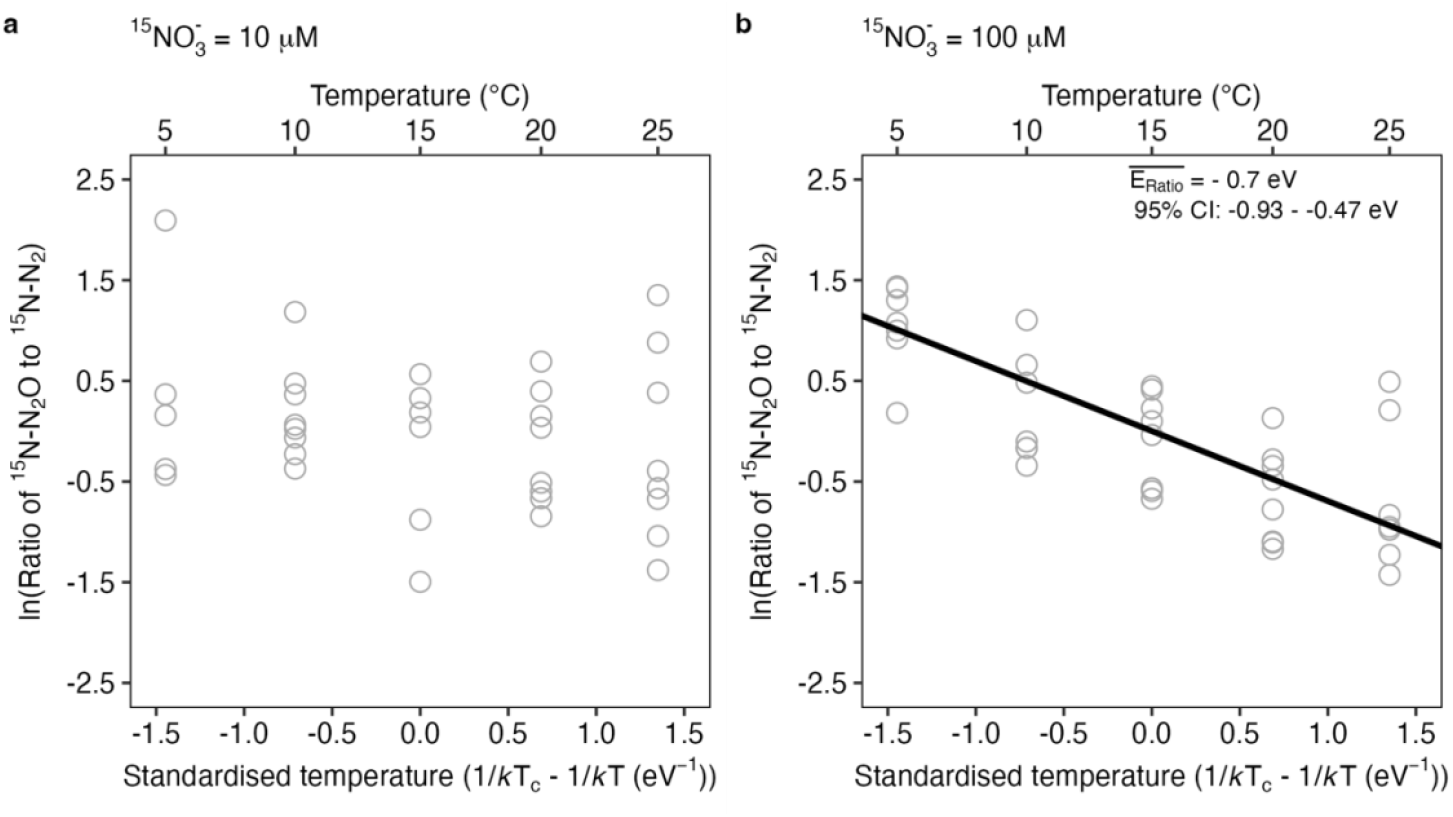
Temperature sensitivity of the ratio of^15^N_2_O and^15^N_2_net production from denitrification under nitrate-limited or nitrate-replete conditions. **a**, Ratio of net production rate of N_2_O and N_2_ was consistent between different temperatures with 10 µM ^15^NO_3_^-^ added. **b**, Ratio of net production rate of N_2_O and N_2_ decreased at higher temperatures with 100 µM of ^15^NO_3_^-^. The temperature was centered at the median temperature of all the data points, i.e. 15°C, while the ratios of N_2_O and N_2_ net production were natural log (ln) transformed and then centered by subtracting the pond-specific intercepts. We visualized the data using the “Visreg” package in R, with the solid line in **b** showing the best-fitting linear mixed-effect model (Supplementary Table 4). The data shown are with the full range of incubation temperatures from 5ºC to 25ºC. *n* = 35 and 38 incubations, respectively, for panels **a** and **b**, from 8 ponds for each ^15^NO_3_^-^ treatment.

## Discussions

From the meta-data analysis of previous studies, both net N_2_O and N_2_ production from denitrification increased at higher temperatures, with apparent activation energies of 0.41 eV and 0.6 eV, respectively (Fig. 1). The temperature response of net N_2_O and N_2_ production appeared consistent across aquatic sediments and terrestrial soils, but available data from aquatic sediments remain sparse. In addition, the N_2_O:N_2_ ratio declined at higher temperatures, reflecting the differing temperature sensitivities of the two gases. However, most of these studies have been carried in N-rich environments or under high NO_3_^-^ enrichment (Table 1), leaving N-limited ecosystems largely unexplored. Moreover, few studies have applied ^15^N-labelling techniques, meaning that N_2_O or N_2_ production have often been attributed to denitrification without excluding contributions from other microbial pathways.

Here, we addressed these limitations by applying ^15^N-labelling techniques to directly isolate denitrification and quantify its temperature sensitivity in N-limited aquatic sediments, an ecosystem type that remains largely overlooked. Our results showed that while both net production and reduction of N_2_O occurred across a temperature range of 5ºC to 25ºC under both 10 and 100 µM ^15^NO_3_^-^, significant temperature effects were only evident at the higher nitrate concentration (Fig. 4**b**, 4**d**). Under nitrate-limited conditions (10 µM ^15^NO_3_^-^), neither net N_2_O production nor its reduction to N_2_ responded significantly to temperature changes (Fig. 4**a**, 4**c**). Similarly, a previous study reported that both the rate and temperature sensitivity of N_2_ production were substantially higher in nitrogen-enriched estuarine mesocosms compared with controls, with activation energies of 1.1 eV and 0.4 eV, respectively (Nowicki, 1994). Together, these results indicate that nitrate availability modulates the temperature sensitivity of N_2_O and N_2_ production from denitrification (Palacin‐Lizarbe et al., 2018).

According to Arrhenius kinetics, enzyme-mediated rates generally increase with temperatures. However, when substrate availability is low, microbial activity can be constrained by substrate supply rather than enzyme kinetics, resulting in minimal temperature sensitivity. Beyond the denitrification patterns observed in this study, such invariant temperature responses under substrate limitation have also been reported for other microbial processes, including N_2_O fixation (Si et al., 2023), ammonia oxidation (Horak et al., 2013; Zheng et al., 2020), and methane oxidation (Lofton et al., 2014; Szafranek-Nakonieczna et al., 2019). Under such conditions, heterotrophic denitrifiers could behave like autotrophs, which often operate under chronic substrate limitation and therefore exhibit little thermal responses.

In the high nitrate treatment (100 µM ^15^NO_3_^-^), N_2_ production increased exponentially from 5 ºC to 25 ºC (Fig. 4**d**). On the contrary, net N_2_O production decreased (Fig. 4**b**), resulting in lower N_2_O accumulation at higher temperatures. This opposing temperature response of N_2_O and N_2_ production has also been reported in river sediments, where rising temperatures led to increased N_2_ production and decreased N_2_O production (Silvennoinen et al., 2008). Similar patterns have been found in soils - with N_2_ production increasing while net N_2_O production declined from 10 ºC to 30 ºC (Bailey, 1976). Elevated N_2_O emissions have also been documented in soils under very low temperatures – for example, at -1 ºC compared to above 0 °C (Wertz et al., 2013), at 0 °C compared to 5 °C (Holtan-Hartwig et al., 2002), and at 4 °C compared to 20 °C (Melin and Nômmik, 1983). Together, these findings indicate that while increasing temperature enhances N_2_ production via complete denitrification, it does not necessarily lead to increased net N_2_O production.

As N_2_ production exhibited greater temperature sensitivity than net N_2_O production in freshwater communities (Fig. 4), the ratio of N_2_O to N_2_ production declined at higher temperatures (Fig. 5). This is consistent with previous studies showing that while N_2_ production from denitrification increases with temperature in both aquatic (Brin et al., 2014; 2017; Nowicki, 1994; Rysgaard et al., 2004; Seitzinger et al., 1984; Silvennoinen et al., 2008; Veraart et al., 2011) and terrestrial ecosystems (Bailey, 1976; Castaldi, 2000; Holtan-Hartwig et al., 2002; Keeney et al., 1979; Lai and Denton, 2018; Qin et al., 2014; Yu et al., 2023), lower product ratios of net N_2_O to N_2_ at higher temperatures were found in studies that have characterised the temperature sensitivity of both net N_2_O to N_2_ in soils (Bailey, 1976; Keeney et al., 1979; Lai and Denton, 2018; Maag and Vinther, 1996) and river sediments (Silvennoinen et al., 2008). Moreover, biogeochemical modelling supports this pattern: for example, the ratio of N_2_O to N_2_ from denitrification was predicted to decrease with warming, leading to a 6% decrease in N_2_O emissions in response to a 1.8°C temperature increase in European forest soils (Kesik et al., 2006).

These findings suggest that N_2_O production and its reduction to N_2_ during denitrification exhibit differing temperature sensitivities. On possible reason is that N_2_O production has a lower activation energy than its reduction to N_2_, resulting in a declining N_2_O:N_2_ production ratio at increasing temperature. Supporting this, incubations of Arctic soils showed that the abundance of *nosZ* - the gene encoding N_2_O reductase - was higher at 10°C than at 4°C, whereas the abundance of *norB*, which is involved in N_2_O production, decreased with rising temperature (Jung et al., 2011). Additional studies in soils have suggested that increased N_2_O emissions at low temperatures may result from the inhibited activity of N_2_O reductase under cold conditions (e.g., around 0ºC) (Holtan-Hartwig et al., 2002; Öquist et al., 2007). Conversely, N_2_O reduction to N_2_ may be enhanced at higher temperatures, potentially due to reduced oxygen solubility or lower oxygen concentrations driven by elevated respiration relative to photosynthesis (Smith, 1997; Veraart et al., 2011).

Our results also showed that both net N_2_O production and the N_2_O: N_2_ ratio decreased with increasing temperature (Fig. 4, Fig. 5), indicating greater relative accumulation of N_2_O at colder temperatures. This additional N_2_O pool could be available for other nitrogen cycling processes, including N_2_O fixation, as suggested by our earlier work showing higher N_2_O fixation:N2 fixation ratios under cold conditions (Si et al., 2023). Such a shift may represent a nitrogen-conserving strategy in nitrogen-limited ecosystems, particularly when N_2_ fixation is energetically less favourable, highlighting a potential link between denitrification and N_2_O fixation in cold environments.

## Conclusions

Here, we characterised the combined effects of temperature and nitrate availability on net N_2_O and N_2_ production from denitrification in understudied N-limited aquatic sediments. Our results showed that the effects of warming on N_2_O and N_2_ production from denitrification are strongly modulated by nitrate availability. In N-limited ecosystems, substrate availability may outweigh temperature in determining the balance between N_2_O to N_2_ production. Under high nitrate concentrations, warming enhanced complete denitrification by reducing fixed nitrogen into N_2_, without necessarily increasing emissions of the atmospherically potent gas N_2_O. These findings highlight the need to consider both nutrient availability and temperature when assessing N_2_O emissions and nitrogen balance in natural waters, as neglecting nutrient limitation may overestimate the impacts of warming.

## Acknowledgements

This study was supported through a PhD Studentship from Queen Mary University of London. We thank M. Rouen for designing and installing the data-logging system for the ponds and J. Pretty for routine maintenance of the ponds.

## Author contributions

M.T and Y.S conceived the study. Y.S performed incubations, analysed the data and wrote the manuscript. Both authors contributed to revisions of the manuscript.

## Competing Interests

The authors declare no competing interests.

## Notes

### Competing Interest Statement

The authors have declared no competing interest.

## References

Bailey, L. 1976. Effects of temperature and root on denitrification in a soil. Canadian Journal of Soil Science 56(2), 79–87.

Bailey, L. and Beauchamp, E. 1973. Effects of temperature on NO_3_^−^ and NO_2_^−^ reduction, nitrogenous gas production, and redox potential in a saturated soil. Canadian Journal of Soil Science 53(2), 213–218.

Bates, D., Mächler, M., Bolker, B. and Walker, S. 2014. Fitting linear mixed-effects models using lme4. arXiv preprint arXiv:1406.5823.

Baulch, H.M., Schiff, S.L., Maranger, R. and Dillon, P.J. 2011. Nitrogen enrichment and the emission of nitrous oxide from streams. Global Biogeochemical Cycles 25(4), GB4013.

Beaulieu, J.J., Smolenski, R.L., Nietch, C.T., Townsend-Small, A., Elovitz, M.S. and Schubauer-Berigan, J.P. 2014. Denitrification alternates between a source and sink of nitrous oxide in the hypolimnion of a thermally stratified reservoir. Limnology and Oceanography 59(2), 495–506.

Beaulieu, J.J., Tank, J.L., Hamilton, S.K., Wollheim, W.M., Hall, R.O., Mulholland, P.J., Peterson, B.J., Ashkenas, L.R., Cooper, L.W., Dahm, C.N., Dodds, W.K., Grimm, N.B., Johnson, S.L., McDowell, W.H., Poole, G.C., Valett, H.M., Arango, C.P., Bernot, M.J., Burgin, A.J., Crenshaw, C.L., Helton, A.M., Johnson, L.T., O’Brien, J.M., Potter, J.D., Sheibley, R.W., Sobota, D.J. and Thomas, S.M. 2011. Nitrous oxide emission from denitrification in stream and river networks. Proceedings of the National Academy of Sciences 108(1), 214–219.

Benoit, M., Garnier, J. and Billen, G. 2015. Temperature dependence of nitrous oxide production of a luvisolic soil in batch experiments. Process Biochemistry 50(1), 79–85.

Breheny, P. and Burchett, W. 2017. Visualization of regression models using visreg. R J. 9(2), 56.

Brin, L.D., Giblin, A.E. and Rich, J.J. 2014. Environmental controls of anammox and denitrification in southern New England estuarine and shelf sediments. Limnology and Oceanography 59(3), 851–860.

Brin, L.D., Giblin, A.E. and Rich, J.J. 2017. Similar temperature responses suggest future climate warming will not alter partitioning between denitrification and anammox in temperate marine sediments. Global change biology 23(1), 331–340.

Castaldi, S. 2000. Responses of nitrous oxide, dinitrogen and carbon dioxide production and oxygen consumption to temperature in forest and agricultural light-textured soils determined by model experiment. Biology and Fertility of Soils 32, 67–72.

Codispoti, L., Brandes, J.A., Christensen, J., Devol, A., Naqvi, S., Paerl, H.W. and Yoshinari, T. 2001. The oceanic fixed nitrogen and nitrous oxide budgets: Moving targets as we enter the anthropocene? Scientia Marina 65(S2), 85–105.

Dalsgaard, T., Canfield, D.E., Petersen, J., Thamdrup, B. and Acuña-González, J. 2003. N2 production by the anammox reaction in the anoxic water column of Golfo Dulce, Costa Rica. Nature 422(6932), 606–608.

Del Prado, A., Merino, P., Estavillo, J., Pinto, M. and González-Murua, C. 2006. N_2_O and NO emissions from different N sources and under a range of soil water contents. Nutrient cycling in agroecosystems 74(3), 229–243.

Dobbie, K. and Smith, K. 2001. The effects of temperature, water‐filled pore space and land use on N_2_O emissions from an imperfectly drained gleysol. European Journal of Soil Science 52(4), 667–673.

Duan, P., Song, Y., Li, S. and Xiong, Z. 2019. Responses of N_2_O production pathways and related functional microbes to temperature across greenhouse vegetable field soils. Geoderma 355, 113904.

Ginestet, P., Audic, J.-M., Urbain, V. and Block, J.-C. 1998. Estimation of nitrifying bacterial activities by measuring oxygen uptake in the presence of the metabolic inhibitors allylthiourea and azide. Applied and Environmental Microbiology 64(6), 2266–2268.

Holtan-Hartwig, L., Dörsch, P. and Bakken, L. 2002. Low temperature control of soil denitrifying communities: kinetics of N_2_O production and reduction. Soil Biology and Biochemistry 34(11), 1797–1806.

Horak, R.E.A., Qin, W., Schauer, A.J., Armbrust, E.V., Ingalls, A.E., Moffett, J.W., Stahl, D.A. and Devol, A.H. 2013. Ammonia oxidation kinetics and temperature sensitivity of a natural marine community dominated by Archaea. The ISME Journal 7(10), 2023–2033.

Ji, Q., Buitenhuis, E., Suntharalingam, P., Sarmiento, J.L. and Ward, B.B. 2018. Global nitrous oxide production determined by oxygen sensitivity of nitrification and denitrification. Global Biogeochemical Cycles 32(12), 1790–1802.

Jung, J., Yeom, J., Kim, J., Han, J., Lim, H.S., Park, H., Hyun, S. and Park, W. 2011. Change in gene abundance in the nitrogen biogeochemical cycle with temperature and nitrogen addition in Antarctic soils. Research in microbiology 162(10), 1018–1026.

Jørgensen, K.S. 1989. Annual pattern of denitrification and nitrate ammonification in estuarine sediment. Applied and Environmental Microbiology 55(7), 1841–1847.

Keeney, D., Fillery, I. and Marx, G. 1979. Effect of temperature on the gaseous nitrogen products of denitrification in a silt loam soil. Soil Science Society of America Journal 43(6), 1124–1128.

Kesik, M., Brüggemann, N., Forkel, R., Kiese, R., Knoche, R., Li, C., Seufert, G., Simpson, D. and Butterbach‐Bahl, K. 2006. Future scenarios of N_2_O and NO emissions from European forest soils. Journal of Geophysical Research: Biogeosciences 111(G2).

Kirkwood, D. 1996. Nutrients: Practical notes on their determination in sea water.

Knowles, R. 1982. Denitrification. Microbiological reviews 46(1), 43.

Kurganova, I. and Lopes de Gerenyu, V. 2010. Effect of the temperature and moisture on the N_2_O emission from some arable soils. Eurasian Soil Science 43, 919–928.

Kuypers, M.M., Marchant, H.K. and Kartal, B. 2018. The microbial nitrogen-cycling network. Nature Reviews Microbiology 16(5), 263.

Kuypers, M.M., Sliekers, A.O., Lavik, G., Schmid, M., Jørgensen, B.B., Kuenen, J.G., Sinninghe Damsté, J.S., Strous, M. and Jetten, M.S. 2003. Anaerobic ammonium oxidation by anammox bacteria in the Black Sea. Nature 422(6932), 608–611.

Lai, T.V. and Denton, M.D. 2018. N_2_O and N_2_ emissions from denitrification respond differently to temperature and nitrogen supply. Journal of Soils and Sediments 18(4), 1548–1557.

Li, Y., Tian, H., Yao, Y., Shi, H., Bian, Z., Shi, Y., Wang, S., Maavara, T., Lauerwald, R. and Pan, S. 2024. Increased nitrous oxide emissions from global lakes and reservoirs since the pre-industrial era. Nature Communications 15(1), 942.

Lofton, D.D., Whalen, S.C. and Hershey, A.E. 2014. Effect of temperature on methane dynamics and evaluation of methane oxidation kinetics in shallow Arctic Alaskan lakes. Hydrobiologia 721, 209–222.

Maag, M. and Vinther, F.P. 1996. Nitrous oxide emission by nitrification and denitrification in different soil types and at different soil moisture contents and temperatures. Applied Soil Ecology 4(1), 5–14.

Maavara, T., Lauerwald, R., Laruelle, G.G., Akbarzadeh, Z., Bouskill, N.J., Van Cappellen, P. and Regnier, P. 2019. Nitrous oxide emissions from inland waters: Are IPCC estimates too high? Global change biology 25(2), 473–488.

Masson-Delmotte, V., Zhai, P., Pirani, A., Connors, S.L., Péan, C., Berger, S., Caud, N., Chen, Y., Goldfarb, L. and Gomis, M. 2021. Climate change 2021: the physical science basis. Contribution of working group I to the sixth assessment report of the intergovernmental panel on climate change 2.

McKenney, D., Johnson, G. and Findlay, W. 1984. Effect of temperature on consecutive denitrification reactions in Brookston clay and Fox sandy loam. Applied and environmental microbiology 47(5), 919–926.

Meinshausen, M., Smith, S.J., Calvin, K., Daniel, J.S., Kainuma, M.L., Lamarque, J.-F., Matsumoto, K., Montzka, S.A., Raper, S.C. and Riahi, K. 2011. The RCP greenhouse gas concentrations and their extensions from 1765 to 2300. Climatic change 109(1), 213–241.

Melin, J. and Nômmik, H. 1983. Denitrification measurements in intact soil cores. Acta Agriculturae Scandinavica 33(2), 145–151.

Myrstener, M., Jonsson, A. and Bergström, A.-K. 2016. The effects of temperature and resource availability on denitrification and relative N_2_O production in boreal lake sediments. Journal of Environmental Sciences 47, 82–90.

Nowicki, B.L. 1994. The effect of temperature, oxygen, salinity, and nutrient enrichment on estuarine denitrification rates measured with a modified nitrogen gas flux technique. Estuarine, Coastal and Shelf Science 38(2), 137–156.

Palacin‐Lizarbe, C., Camarero, L. and Catalan, J. 2018. Denitrification Temperature Dependence in Remote, Cold, and N‐Poor Lake Sediments. Water Resources Research 54(2), 1161–1173.

Qin, S., Yuan, H., Hu, C., Oenema, O., Zhang, Y. and Li, X. 2014. Determination of potential N_2_O-reductase activity in soil. Soil Biology and Biochemistry 70, 205–210.

Ravishankara, A., Daniel, J.S. and Portmann, R.W. 2009. Nitrous oxide (N_2_O): the dominant ozone-depleting substance emitted in the 21st century. Science 326(5949), 123–125.

Rysgaard, S., Glud, R.N., Risgaard-Petersen, N. and Dalsgaard, T. 2004. Denitrification and anammox activity in Arctic marine sediments. Limnology and oceanography 49(5), 1493–1502.

Seitzinger, S.P., Nixon, S.W. and Pilson, M.E. 1984. Denitrification and nitrous oxide production in a coastal marine ecosystem 1. Limnology and Oceanography 29(1), 73–83.

Si, Y., Zhu, Y., Sanders, I., Kinkel, D.B., Purdy, K.J. and Trimmer, M. 2023. Direct biological fixation provides a freshwater sink for N_2_O. Nature Communications 14(1), 6775.

Silvennoinen, H., Liikanen, A., Torssonen, J., Stange, C. and Martikainen, P. 2008. Denitrification and N_2_O effluxes in the Bothnian Bay (northern Baltic Sea) river sediments as affected by temperature under different oxygen concentrations. Biogeochemistry 88, 63–72.

Smith, K. 1997. The potential for feedback effects induced by global warming on emissions of nitrous oxide by soils. Global Change Biology 3(4), 327–338.

Smith, K., Thomson, P., Clayton, H., McTaggart, I. and Conen, F. 1998. Effects of temperature, water content and nitrogen fertilisation on emissions of nitrous oxide by soils. Atmospheric Environment 32(19), 3301–3309.

Szafranek-Nakonieczna, A., Wolińska, A., Zielenkiewicz, U., Kowalczyk, A., Stępniewska, Z. and Błaszczyk, M. 2019. Activity and identification of methanotrophic bacteria in arable and no-tillage soils from Lublin region (Poland). Microbial Ecology 77(3), 701–712.

Team, R.C. 2021. R: A language and environment for statistical computing.

Trimmer, M., Engstrom, P. and Thamdrup, B. 2013. Stark contrast in denitrification and anammox across the deep Norwegian trench in the Skagerrak. Appl Environ Microbiol 79(23), 7381–7389.

Trimmer, M., Risgaard-Petersen, N., Nicholls, J.C. and Engström, P. 2006. Direct measurement of anaerobic ammonium oxidation (anammox) and denitrification in intact sediment cores. Marine Ecology Progress Series 326, 37–47.

Veraart, A.J., De Klein, J.J. and Scheffer, M. 2011. Warming can boost denitrification disproportionately due to altered oxygen dynamics. PloS one 6(3), e18508.

Warren, V. (2017) The temperature dependence of the gaseous products of the nitrogen cycle, Queen Mary University of London.

Weiss, R. and Price, B. 1980. Nitrous oxide solubility in water and seawater. Marine chemistry 8(4), 347–359.

Weiss, R.F. 1970 The solubility of nitrogen, oxygen and argon in water and seawater, pp. 721–735, Elsevier.

Wertz, S., Goyer, C., Zebarth, B.J., Burton, D.L., Tatti, E., Chantigny, M.H. and Filion, M. 2013. Effects of temperatures near the freezing point on N_2_O emissions, denitrification and on the abundance and structure of nitrifying and denitrifying soil communities. FEMS microbiology ecology 83(1), 242–254.

Yu, H., Duan, Y., Mulder, J., Dörsch, P., Zhu, W., Huang, K., Zheng, Z., Kang, R., Wang, C. and Quan, Z. 2023. Universal temperature sensitivity of denitrification nitrogen losses in forest soils. Nature Climate Change 13(7), 726–734.

Yvon-Durocher, G., Allen, A.P., Bastviken, D., Conrad, R., Gudasz, C., St-Pierre, A., Thanh-Duc, N. and Del Giorgio, P.A. 2014. Methane fluxes show consistent temperature dependence across microbial to ecosystem scales. Nature 507(7493), 488.

Zheng, Z.-Z., Zheng, L.-W., Xu, M.N., Tan, E., Hutchins, D.A., Deng, W., Zhang, Y., Shi, D., Dai, M. and Kao, S.-J. 2020. Substrate regulation leads to differential responses of microbial ammonia-oxidizing communities to ocean warming. Nature Communications 11(1), 3511.

Zhu, G., Shi, H., Zhong, L., He, G., Wang, B., Shan, J., Han, P., Liu, T., Wang, S. and Liu, C. 2025. Nitrous oxide sources, mechanisms and mitigation. Nature Reviews Earth & Environment, 1–19.

Zhu, Y., Purdy, K.J., Eyice, Ö., Shen, L., Harpenslager, S.F., Yvon-Durocher, G., Dumbrell, A.J. and Trimmer, M. 2020. Disproportionate increase in freshwater methane emissions induced by experimental warming. Nature Climate Change, 1–6.

Öquist, M.G., Petrone, K., Nilsson, M. and Klemedtsson, L. 2007. Nitrification controls N_2_O production rates in a frozen boreal forest soil. Soil Biology and Biochemistry 39(7), 1809–1811.

