## Supplementary information for "Nitrate availability modulates the temperature effect on N_2_O and N_2_ production from denitrification"

**Supplementary Figure 1 | The $^{15}$N-labelling of the $NO_3^-$ pool being denitrified ($F_N$, calculated using the $^{15}$N-labelling of $N_2$ produced in the incubations) in incubations with different concentrations of $^{15}NO_3^-$ added.** $F_N$ was lower in experiments with less $^{15}NO_3^-$ added, indicating the interference from background $^{14}NO_x^-$ that has a stronger effect on the $^{15}$N-labelling at lower $^{15}NO_3^-$ concentrations. The dots are the means, with vertical bars showing the standard error (*n* = 6 ponds for each concentration of $^{15}NO_3^-$).

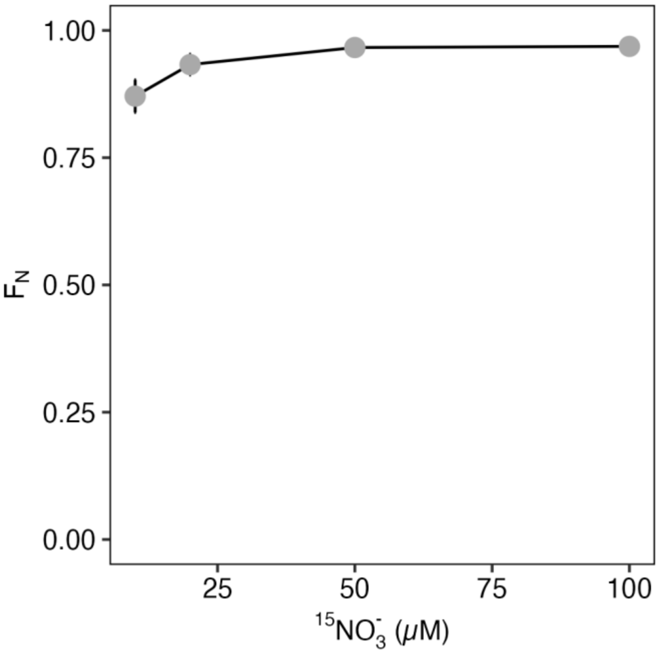

**Supplementary Figure 2 | The accumulation of $^{15}N_2O$ and $^{15}N_2$ over time from independent sediment incubations amended with different concentrations of $^{15}NO_3^-$. a**, $^{15}N_2O$ production quickly reached a peak and started to decline in incubations with $^{15}NO_3^-$ additions below 100 μM, while with 100 μM of $^{15}NO_3^-$, the peak occurred later, at approximately 3 hours. **b**, $^{15}N_2$ continued to accumulate over the 24-hour incubation period with $^{15}NO_3^-$ additions above 10 μM, while with 10 μM of $^{15}NO_3^-$, it plateaued at approximately 12 hours. The dots in each plot represent the means, with vertical bars showing the standard error (*n* = 6 ponds for each concentration of $^{15}NO_3^-$ at each time point).

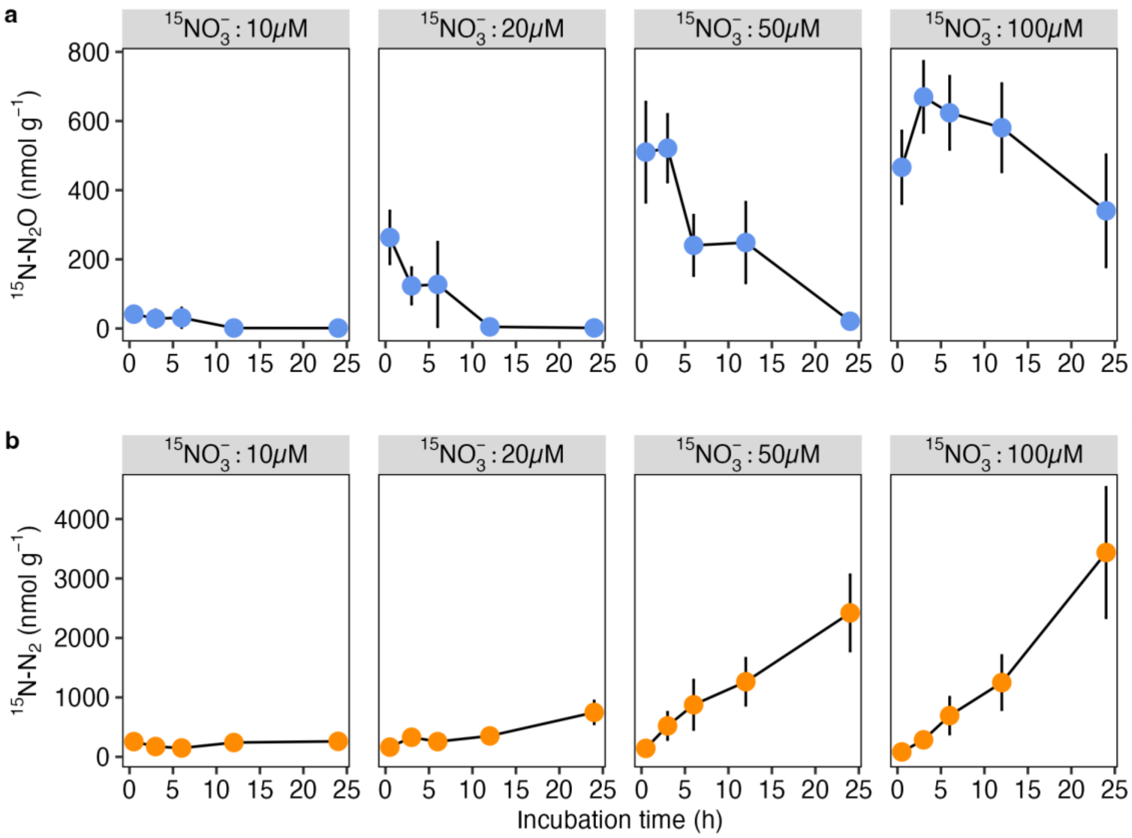

**Supplementary Figure 3 | Excess $^{29}N_2$ in the treatments compared to controls from the experiments to confirm whether anammox activity is present in the experimental ponds.** No additional $^{29}N_2$ was produced from $^{15}NH_4^+$ plus $^{14}NO_3^-$ treatments relative to the $^{15}NH_4^+$ only treatments, which indicated that anammox was not present in the ponds and that denitrification could be treated as the only $N_2$ producing process.

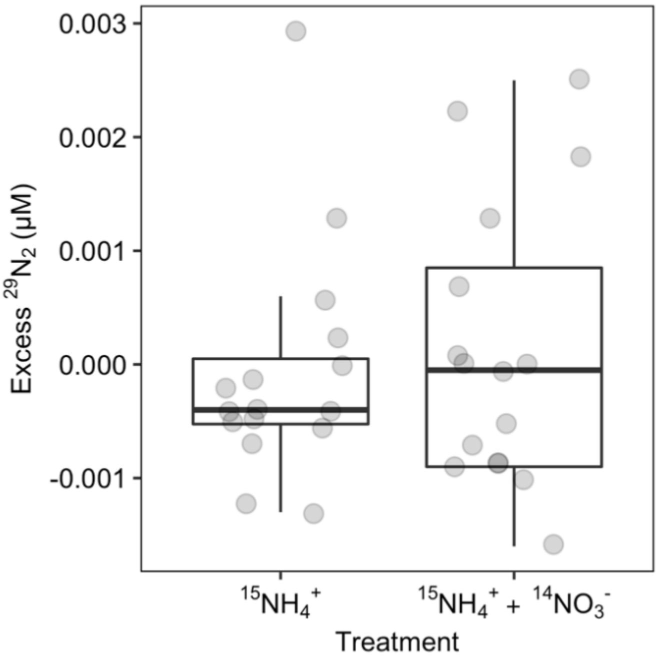

**Supplementary Figure 4 |** No significant net $^{15}N-N_2O$ production was detected in sediments from most ponds. Net $^{15}N-N_2O$ production via nitrification was observed in only one of eight pond: it occurred at all time points with 44 μM $^{15}NH_4^+$, at 3 h with 22 μM of $^{15}NH_4^+$, and was absent when nitrification was inhibited with ATU.

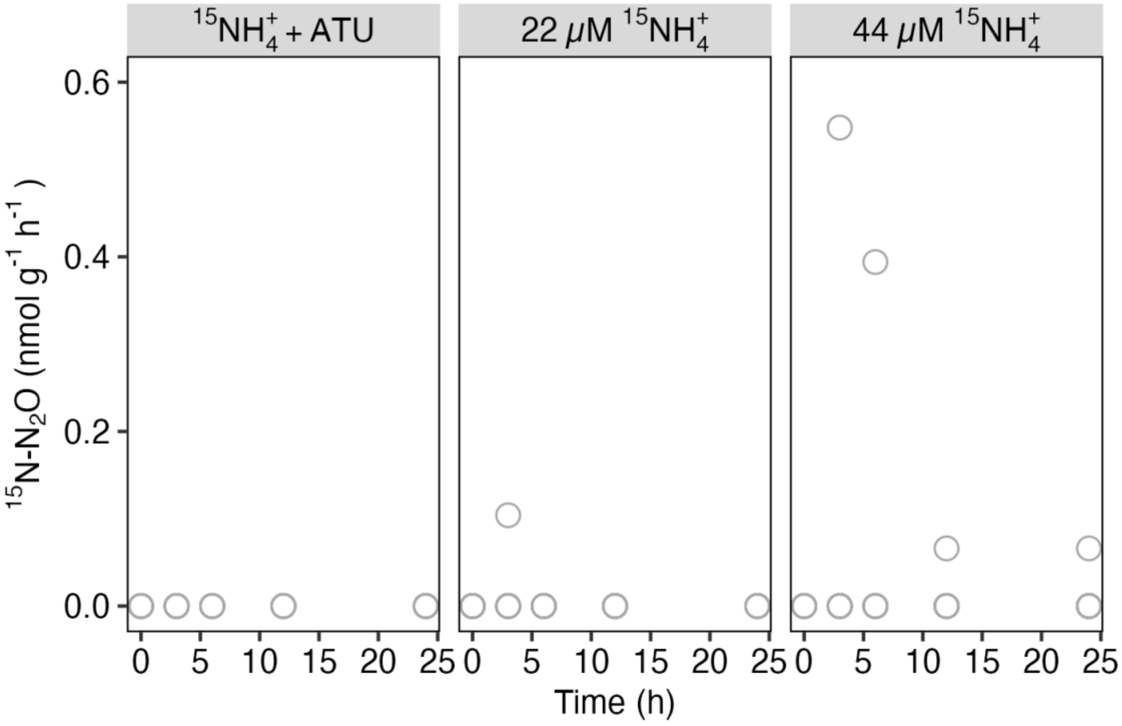

**Supplementary Figure 5 | After 24 hours of incubation, the ratio (%) of $^{15}N$-$N_2$ production to $^{15}N$-$NO_3^-$ addition was consistent between different $^{15}NO_3^-$ concentrations (42.9% ± 3.1%, mean ± s.e.; *p* = 0.52), indicating that the fraction of substrate denitrified was not affected by the availability of $NO_3^-$.** The percentage of $^{15}N$-$N_2$ produced to $^{15}NO_3^-$ added in the incubations was calculated by $({}^{29}N_2 + 2\times{}^{30}N_2) / {}^{15}NO_3^- \times 100$ (all in nmol N $g^{-1}$ sediment), where the difference in the fraction between incubations with different $^{15}NO_3^-$ additions was tested by the Kruskal-Wallis test.

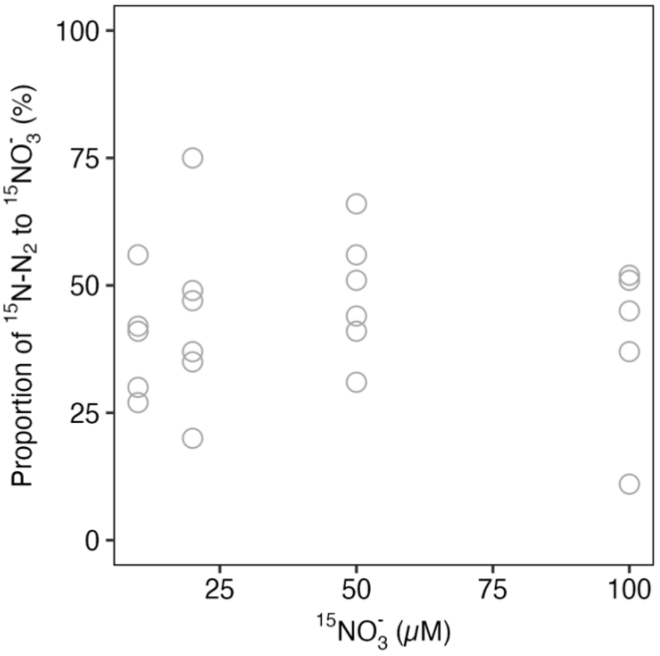

**Supplementary Table 1 | Concentration of porewater nutrients and the $^{15}N$-labelling of the $NH_4^+$ pool ($F_A$) with 44 μM of $^{15}NH_4^+$ added at the start of the nitrification experiments.** NA: not available. Mean: mean value of each category from all ponds. s.e.: standard error.

| Pond | $NO_2^-$ | $NO_3^-$ | $NH_4^+$ | $PO_4^{3-}$ | $F_A$ |
|---|---|---|---|---|---|
| 2 | 0.31 | 0.77 | 3.78 | 0.57 | 0.89 |
| 5 | NA | NA | NA | 0.44 | 0.98 |
| 7 | 0.20 | 0.2 | 1.84 | 0.88 | 0.95 |
| 8 | 0.06 | 0.18 | 0.98 | 0.61 | 0.96 |
| 9 | 0.18 | 0.16 | 66.3 | 1.39 | 0.41 |
| 11 | 0.43 | 0.7 | 3.71 | 0.60 | 0.90 |
| 12 | 0.60 | 1.78 | 10.9 | 0.84 | 0.78 |
| 17 | 0.18 | 0.26 | 1.35 | 0.82 | 0.95 |
| Mean | 0.28 | 0.58 | 12.7 | 0.77 | 0.85 |
| s.e. | 0.07 | 0.22 | 9.03 | 0.10 | 0.07 |

**Supplementary Table 2 | Meta-analysis for published rates of total $N_2O$, $N_2$ and net $N_2O$ from denitrification as a function of incubation temperature from both aquatic sediments and terrestrial soils (Fig. 1).** Linear mixed-effects model selection included centred temperature (Tc) and the interaction between Tc and ecosystem type as fixed effects, with a random intercept (1|Study) and slope (0+Tc|Study) to account for variation across the different studies (Table 1). Here, the best-fitting models (**M0**) showed that the rate of net $N_2O$ and $N_2$ production both increased at higher temperatures, and that their temperature sensitivities were not different between aquatic and terrestrial ecosystems. Models were ranked by the small sample-size corrected Akaike Information Criterion (AICc) with the best models (in **bold**) having the lowest AIC values. The null model (M3 for $N_2O$ and M4 for $N_2$) only included an intercept, denoted by 1. lnNet.$N_2O$ and ln$N_2$ are the natural log (ln) transformed rate of net $N_2O$ and $N_2$ production, respectively, and Ecosystem is ecosystem type (terrestrial or aquatic). Each model was compared to the best model using the Log-likelihood ratio test (LogLik, d.f. degrees of freedom) showing $\chi^2$ (Chi-squared statistic) and *p* (the corresponding *p*-value).

| Model | d.f. | AICc | LogLik | $\chi^2$ | *p* |
|---|---|---|---|---|---|
| **Net $N_2O$ production** | | | | | |
| **M0:** lnNet.$N_2O$~Tc | 4 | 959.7 | -475.8 | | |
| M1: lnNet.$N_2O$~Tc+Ecosystem | 5 | 961.4 | -475.6 | 0.39 | 0.53 |
| M2: lnNet.$N_2O$~Tc*Ecosystem | 6 | 963.3 | -475.5 | 0.61 | 0.77 |
| M3: lnNet.$N_2O$~1 | 3 | 996.8 | -495.4 | 39.2 | **<0.001** |
| M4: lnNet.$N_2O$~Ecosystem | 4 | 998.3 | -495.1 | 38.7 | **<0.001** |
| **$N_2$ production** | | | | | |
| **M0: ln$N_2$~Tc** | 5 | 1288.5 | -639.2 | | |
| M1: ln$N_2$~Tc+Ecosystem | 6 | 1290.5 | -639.2 | 0 | 0.997 |
| M2: ln$N_2$~Tc*Ecosystem | 7 | 1292.5 | -639.1 | 0.11 | 0.95 |
| M3: ln$N_2$~Ecosystem | 4 | 1312.1 | -652.0 | 25.7 | **<0.001** |
| M4: ln$N_2$~1 | 5 | 1314.1 | -652.0 | 25.7 | **<0.001** |
| **$N_2O/N_2$** | | | | | |
| **M0: ln$N_2O/N_2$~Tc** | 4 | 691.2 | -341.5 | | |
| M1: ln$N_2O/N_2$~Tc+Ecosystem | 5 | 693.3 | -341.5 | 0 | 0.99 |
| M2: ln$N_2O/N_2$~Tc*Ecosystem | 6 | 694.0 | -340.8 | 1.44 | 0.49 |
| M3: ln$N_2O/N_2$~Ecosystem | 3 | 702.8 | -348.3 | 13.6 | **<0.001** |
| M4: ln$N_2O/N_2$~1 | 4 | 704.8 | -348.3 | 13.6 | **<0.001** |

**Supplementary Table 3 | Concentrations of porewater nutrients (µM) in sediments used to characterise the temperature sensitivities of $N_2O$ and $N_2$ production from denitrification.** s.e.: standard error (*n* = 8 ponds for each nutrient species).

| Pond | $NO_2^-$ | $NO_3^-$ | $NO_x^-$ | $NH_4^+$ | $PO_4^{3-}$ |
|---|---|---|---|---|---|
| 3 | 0.37 | 1.19 | 1.56 | 5.11 | 0.24 |
| 4 | 0.16 | 0.53 | 0.69 | 41.16 | 0.24 |
| 5 | 0.15 | 0.52 | 0.67 | 1.03 | 0.15 |
| 6 | 0.12 | 0.26 | 0.38 | 3.61 | 0.22 |
| 9 | 0.08 | 0.25 | 0.34 | 31.84 | 0.27 |
| 10 | 0.10 | 0.30 | 0.40 | 25.72 | 0.20 |
| 11 | 0.08 | 0.19 | 0.27 | 0.99 | 0.19 |
| 12 | 0.21 | 0.36 | 0.57 | 0.36 | 0.21 |
| Mean | 0.16 | 0.45 | 0.61 | 13.73 | 0.22 |
| s.e. | 0.03 | 0.11 | 0.15 | 5.83 | 0.01 |

**Supplementary Table 4 | Net production rate of $N_2O$ and $N_2$ (Fig. 4) and their ratio (Fig. 5) as a function of incubation temperature at 10 μM or 100 μM of $NO_3^-$ with sediments from the experimental ponds.** Linear mixed-effects model selection included centered temperature (Tc) as fixed effects, with a random intercept and slope (1|Pond) to account for variation across the different ponds. Separate models were fitted for 10 μM and 100 μM of $NO_3^-$, which were denoted as M10 and M100, respectively. Models were ranked by the small sample-size corrected Akaike Information Criterion (AICc) with the better models (in **bold**) having lower AIC values. Null models only have an intercept, denoted by 1. ln$PN_2O$ and ln$PN_2$ is the natural log (ln) transformed rate of net production of $N_2O$ and $N_2$, respectively, while lnRatio is the ratio of net production of $N_2O$ and $N_2$. Each model was compared to the best model (in **bold**) using the Log-likelihood ratio test (LogLik, d.f. degrees of freedom) showing $\chi^2$ (Chi-squared statistic) and *p* (the corresponding *p*-value).

| **Model** | **d.f.** | **AICc** | **LogLik** | **$\chi^2$** | ***p*** |
|---|---|---|---|---|---|
| **$N_2O$ net production** | | | | | |
| **M10.a: ln$PN_2O$~1** | **3** | **101.3** | **-47.3** | | |
| M10.b: ln$PN_2O$~Tc | 4 | 103.1 | -46.9 | 0.78 | 0.38 |
| **M100.a: ln$PN_2O$~Tc** | **4** | **81.2** | **-36.0** | | |
| M100.b: ln$PN_2O$~1 | 3 | 90.6 | -41.9 | 11.9 | <0.001 |
| **$N_2$ production** | | | | | |
| **M10.a: ln$PN_2$~1** | **3** | **103.1** | **-48.2** | | |
| M10.b: ln$PN_2$~Tc | 4 | 103.8 | -47.3 | 1.79 | 0.18 |
| **M100.a: ln$PN_2$~Tc** | **4** | **109.2** | **-50.0** | | |
| M100.b: ln$PN_2$~1 | 3 | 120.6 | -56.9 | 13.9 | <0.001 |
| **Ratio of $N_2O$ to $N_2$** | | | | | |
| **M10.a: lnRatio~1** | **3** | **106.2** | **-49.7** | | |
| M10.b: lnRatio~Tc | 4 | 106.8 | -48.8 | 1.87 | 0.17 |
| **M100.a: lnRatio~Tc** | **4** | **119.1** | **-54.9** | | |
| M100.b: lnRatio~1 | 3 | 141.1 | -67.2 | 24.6 | <0.001 |
